# Plant cell-based 4D engineered living materials

**DOI:** 10.64898/2026.10.19.760083

**Authors:** Erica Colaprico, Chiara Musumeci, Léa Bogdziewiez, Denise Pagliara, Ariel García Fleitas, Gwennaël Dufil, Elli Magkouta, Xu Jin, Daniele Mantione, Åsa Strand, Stéphane Verger, Eleni Stavrinidou

## Abstract

Engineered living materials (ELMs) seek to bridge the gap between static engineered matter and dynamic biological systems. Photosynthetic ELMs have emerged as a promising route towards autonomous and sustainable material systems; however, they remain largely based on microorganisms or isolated organelles, whereas the potential of living plant cells as multifunctional, adaptive building blocks remains largely unexplored. Here we establish plant cells as a distinct chassis for engineered living materials and demonstrate that their intrinsic metabolic plasticity, cell-wall biochemistry and mechanics can be harnessed to create adaptive, evolvable and programmable materials. Using Arabidopsis thaliana suspension cells and printable pectin–alginate bioinks, we fabricate two- and three-dimensional living materials that support growth, metabolic plasticity and long-term viability. Non-photosynthetic ELMs undergo light-induced metabolic reprogramming to acquire photosynthetic activity, demonstrating adaptive behaviour, while continuous cell growth dynamically remodels the mechanical properties of the material, enabling evolution of material properties through intrinsic biological processes. We further exploit endogenous cell-wall peroxidases to catalyse the biofabrication of conducting polymers, enabling programmable acquisition of electrochemical functionality without genetic modification. By integrating living plant cells with auxetic and honeycomb architectures, we show that biological activity and structural design jointly determine emergent mechanical behaviour, revealing a principle of geometry-dependent biological reinforcement for living mechanical metamaterials. Centimetre-scale plant ELMs retain robust photosynthetic activity, with electron transport rates approaching those of native Arabidopsis tissues despite the absence of leaf architecture. Together, these findings establish plant cells as versatile building blocks for multifunctional living materials and lay the foundation for self-powered autonomous materials with potential applications in soft robotics, bioelectronics and carbon capture.

## Introduction

Biological tissues continuously grow, remodel their structure, adapt their metabolism and evolve their functions in response to physiological and environmental cues. In contrast, most engineered materials are designed with fixed structures and predefined properties that progressively degrade over time. Bridging this divide between inert materials and living adaptive systems has emerged as a central challenge in materials science.

Engineered living materials (ELMs) have emerged at the convergence of materials science, synthetic biology and biofabrication as a new class of materials that integrate living cells within structural matrices to generate dynamic and functional systems^1^. ELMs aim to endow materials with life-like capabilities including growth, self-healing, environmental responsiveness and adaptation. Advances in synthetic biology have enabled the programming of living cells with genetic circuits that sense, process and respond to environmental stimuli^2,3^ while developments in hydrogel engineering and additive manufacturing have provided structurally defined, cell-supportive matrices for integrating biological function into macroscopic materials^4^. To date, ELMs have been developed using bacteria^5,6^, fungi^7,8^, yeast^9^, algae^10,11^ and mammalian cells^12^ for applications including biosensing^5^, bioremediation^13^, soft robotics^12^, therapeutic delivery^14^, energy conversion^15^, and sustainable manufacturing^16,17^.

Photosynthetic ELMs have recently emerged as a promising route towards autonomous and sustainable material systems. Many examples rely on the incorporation of living algae or cyanobacteria within printable hydrogels to create photosynthetically active materials for tissue oxygenation^18^, biologically driven material synthesis^16^, carbon sequestration^11^ and biomineralization^17^. Isolated chloroplasts have also been integrated into synthetic matrices to generate photosynthetically active materials capable of glucose driven growth, self-strengthening and self-repair^19,20^. Despite these advances, current photosynthetic ELMs remain predominantly based on unicellular photosynthetic organisms or isolated organelles. By contrast, the potential of living plant cells, which combine photosynthetic capability with developmental plasticity, cell-wall biomanufacturing and metabolic adaptability, remains largely unexplored as a route towards multifunctional and adaptive living materials.

Recently, the first plant engineered living material was reported, in which genetically transformable tobacco BY-2 cells were bioprinted to create programmable materials with defined pigmentation^21^. Although this represented an important step towards plant-based ELMs, the system relied on non-photosynthetic suspension cells and relatively simple genetic programming. Parallel advances in plant biofabrication have demonstrated the feasibility of bioprinting plant cells within hydrogel matrices for applications ranging from cultured plant tissues for food manufacturing to studies of cellular reprogramming and development^22–24^. However, these efforts have primarily focused on tissue culture and cell characterization rather than on creating living materials whose properties dynamically change through cellular activity.

Here we introduce 4D plant cell–based engineered living materials, where fourth dimension is time, that combine adaptation, evolution and programmability within a single living platform (Fig. 1A). Using *Arabidopsis thaliana* cells and printable hydrogel bioinks, we demonstrate living materials that undergo light-induced metabolic reprogramming, continuously remodel their mechanical properties through growth, and acquire new electrochemical functionality through endogenous biocatalysis. By integrating plant cells with architected geometries, we further show that biological activity and structural design together give rise to emergent mechanical behavior. Collectively, our results establish plant cells as versatile building blocks for a new generation of multifunctional living materials that couple environmental responsiveness, biological growth and biomanufacturing with macroscopic material function.

**Figure 1:**
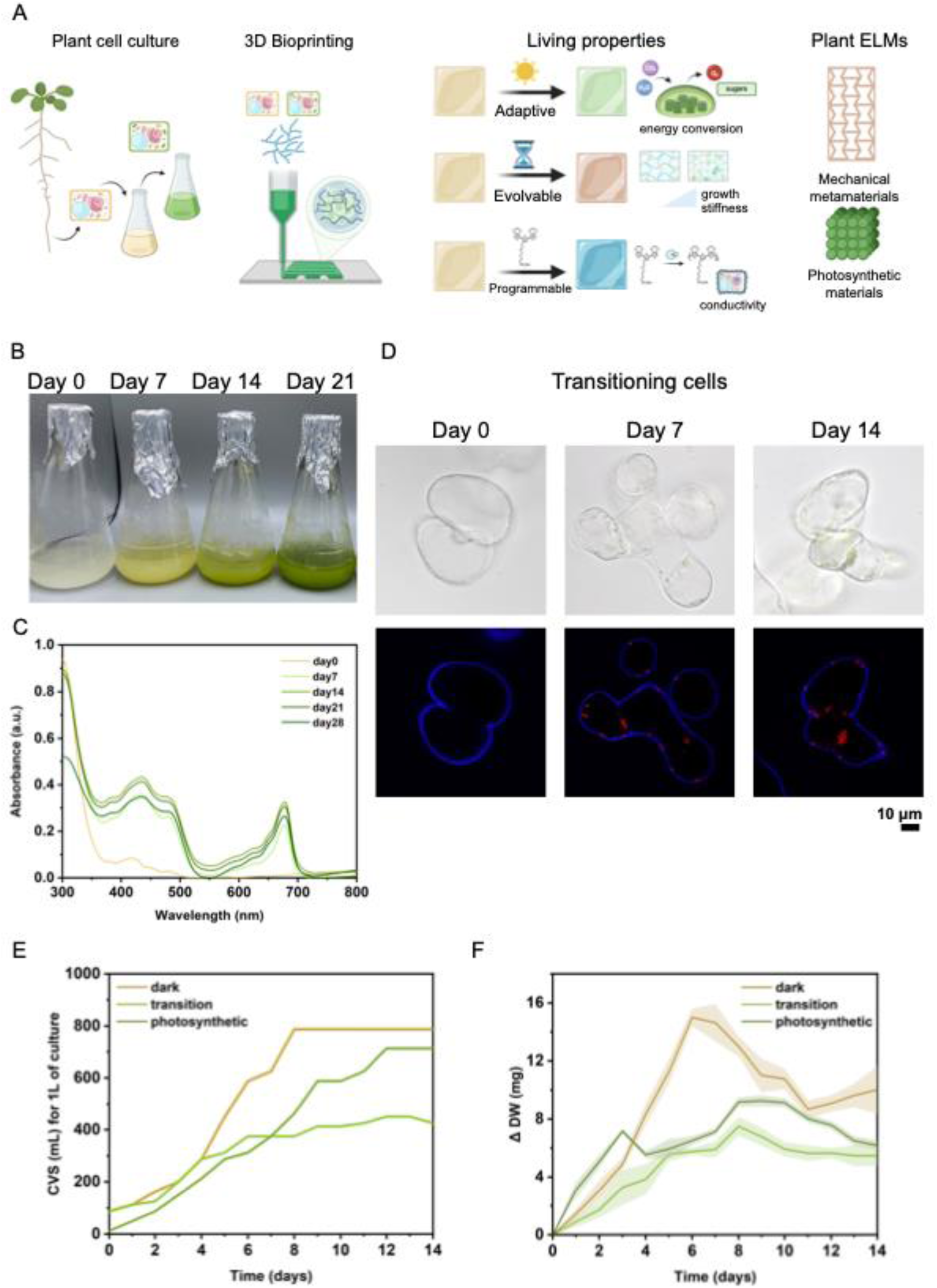
*Arabidopsis thaliana* cells as building blocks of engineered living materials. A. Concept of plant cell–based 4D engineered living materials (ELMs). Photosynthetic and non-photosynthetic Arabidopsis suspension cell cultures are combined with biopolymers to formulate printable bioinks, enabling the fabrication of 2D and 3D plant ELMs. The resulting living materials exhibit adaptive, evolvable and programmable functionalities and are demonstrated as living mechanical metamaterials and photosynthetic materials. B. Transition of plant cells from a non-photosynthetic to a photosynthetic state in culture. Cell suspension culture grown in dark is transferred to growth in light conditions for 21 days, progressively acquiring a green phenotype indicative of photosynthetic activity. C. Absorption spectra of cells during the transition. Characteristic peaks corresponding to chlorophyll a and chlorophyll b are observed after 7 days of light exposure in blue and red region of the spectrum. D. E. Growth curves of non-photosynthetic (denoted dark), transitioning, and photosynthetic cells measured by (D) cell volume after sedimentation (CVS) in mL per litre of suspension culture and (E) dry weight change (ΔDW) (n = 9, mean and standard deviation). F. Representative brightfield and confocal fluorescence micrographs of non-photosynthetic cells (day 0) transitioning to photosynthetic state, (day 7 and day 14) after placing the subculture under light conditions at day 0. Chloroplasts are visible in brightfield images in green. In fluorescence mode, chloroplasts appear red due to chlorophyll autofluorescence while the cell wall appears blue due to staining with calcofluor white. Scale bar, 10 μm.

## Results

### Plant cells enable dynamic, functional building blocks for engineered living materials

To establish plant cells versatility as components of engineered living materials, we first investigated the growth dynamics of suspension cultures of *Arabidopsis thaliana* root cells and their capacity to transition between metabolic states and acquire photosynthetic functionality (Fig. 1B). Cells cultured under dark conditions exhibited a non-photosynthetic phenotype, which progressively shifted upon light exposure, as reported previously^25^. Over a period of 21 days, cells developed a progressively pronounced green coloration, indicating chloroplast development and activation of photosynthesis (Fig. 1B). This transition was confirmed spectroscopically, with the appearance of characteristic chlorophyll a and chlorophyll b absorption peaks in blue and red region of the spectrum, after 7 days of light exposure (Fig. 1C, Fig S1A). Apart from non-photosynthetic and transitioning culture we also established a photosynthetic culture that maintains photosynthetic phenotype over subculturing (Fig S1B).

Growth analysis revealed broadly similar growth trends across metabolic states, with some distinct differences in growth dynamics. Non photosynthetic (denoted dark) cultures exhibited a typical sigmoidal growth profile, characterized by an initial lag phase, accelerated biomass accumulation, and stationary phase by day 8. Photosynthetic cultures showed a transient slowdown around day 6, followed by renewed growth and then stationary phase at day 12, likely associated with media exchange at day 7. In contrast, transitioning cultures, which were transferred from dark to light conditions and in minimal sucrose concentration medium at the start of subculture (day 0), did not exhibit growth acceleration after media replacement and instead reached the stationary phase after day 6 (Fig. 1E).

Dry weight measurements further highlighted differences between metabolic states (Fig. 1F). Biomass accumulation peaked at days 6–7 for non-photosynthetic cells and at days 8–10 for photosynthetic cells. The media-induced enhancement in growth was also reflected in the dry weight increase of photosynthetic cultures. In contrast, transitioning cultures reached peak growth at day 8 and then saturation, suggesting that cellular resources were preferentially allocated towards formation of the photosynthetic apparatus rather than biomass accumulation. Although photosynthetic cells exhibited greater biomass accumulation during the early stages of culture, non-photosynthetic cultures subsequently surpassed them before both converged to similar biomass levels by day 10.

Overall, these results demonstrate that metabolic reprogramming does not highly compromise biomass accumulation (Fig. 1E). Microscopy further confirmed the photosynthetic transition. Chloroplasts were clearly visible in both brightfield and confocal fluorescence images at days 7 and 14 of culture, with chlorophyll autofluorescence observed in red. Cell walls remained intact throughout the transition, as confirmed by calcofluor staining (blue), indicating preserved cellular integrity (Fig. 1D). Together, these findings demonstrate that plant cells can dynamically reconfigure their metabolic state while maintaining viability and functionality, supporting their use as versatile and adaptive building blocks for 4D engineered living materials.

### Bioprinting enables spatially controlled plant-based living materials

We next evaluated the ability of plant cells to form 2D and 3D structured materials through extrusion bioprinting (Fig. 2). To formulate the bioinks, we selected low-methoxy (LM) pectin as a primary component because it acts as the adhesive substance between plant cells and therefore mimics their native extracellular environment. Alginate, although absent from plant cell walls and instead a major component of algal cell walls, was incorporated because of its ability to form stable hydrogels through Ca^2+^-mediated ionic crosslinking via the well-established “egg-box” model^26–28^. Alginate is a linear copolymer of β-D-mannuronic acid and α-L-guluronic acid residues, whose abundant carboxyl groups promote hydrogel formation through a high density of ionic interactions. In contrast, the methyl ester groups present in pectin reduce chain–chain interactions and introduce heterogeneity into the gel network. Combining the two polysaccharides resulted in enhanced gel formation through complementary ionic and hydrogen-bonding interactions between the hydroxyl and carboxyl groups, as well as with the oxygen atoms of the C-O covalent bond of the methoxyl groups (Fig. 2A). Optimization of the pectin– alginate bioink composition (denoted as pec(wt/v%)alg(wt/v%)) showed that increasing the concentration of LM-pectin improved print fidelity, enabling the fabrication of millimetre-thick (∼ 1.5 mm) 2D mesh structures. To fabricate centimetre-scale 3D architectures with high shape fidelity, alginate was replaced with a higher-viscosity grade (denoted alg*) (Fig. 2B, Fig S2).

**Figure 2:**
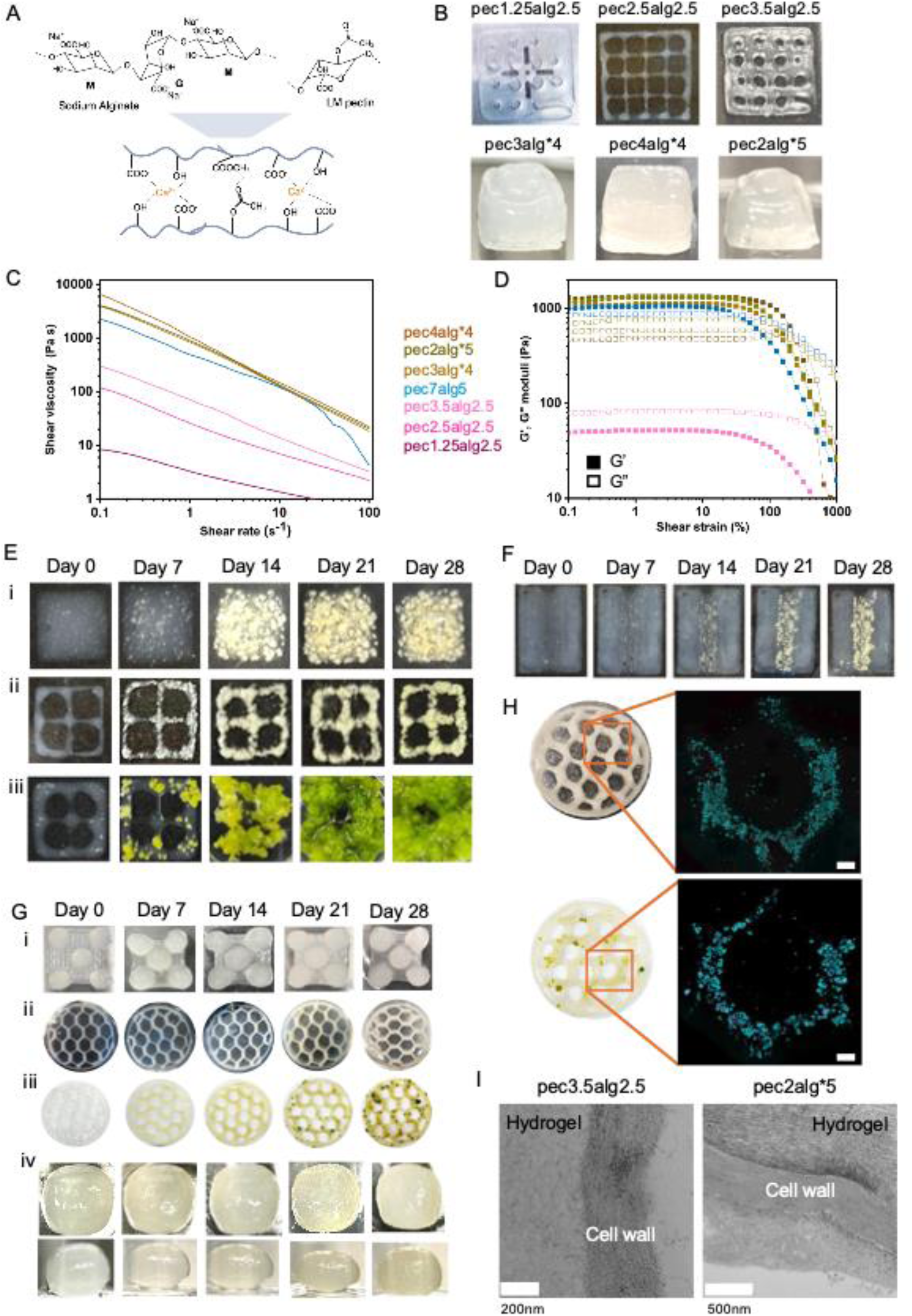
Bioprinted plant cells form 2D and 3D living materials. A. Simplified schematic of intermolecular network between sodium alginate and LM pectin biopolymers’ chains. The crosslinking mechanism relies on the formation of junction zones by Ca^2+^ ions with the negatively charged groups of the two polysaccharides. B. Optimization of pectin-alginate bioink formulations (given in wt/v%) for extrusion bioprinting of 2D and 3D ELMs. Increasing concentrations of low-methoxy (LM) pectin improve shape fidelity in 2x2 cm square mesh structures. For 3D constructs, 1 cm^3^ cubes are printed using high-viscosity alginate (denoted alg*) in combination with LM pectin. C, D. Rheological characterization of selected inks. (C) Shear viscosity as a function of shear rate demonstrates shear-thinning behaviour suitable for extrusion bioprinting. (D) Oscillatory strain sweeps show increased storage (G′) and loss (G″) moduli in inks with high-viscosity alginate and overall polymer content. E. Formation of 2D plant cell–based living materials. Representative photographs over 28 days of growth. i-ii. Non-photosynthetic cells proliferate in full and mesh constructs printed with pec3.5alg2.5 bioink over 28 days maintaining the shape of the construct. iii. Photosynthetic cells printed with pec3.5alg2.5 bioink exhibit less controlled growth in mesh structures over the same period, overgrowing the printed structure. Construct size, 1x1 cm. F. Spatial control of cell growth via matrix composition. Representative photographs over 28 days of growth. Constructs printed with abiotic pec7alg5 in the outer regions and pec3.5alg2.5 bioink in the inner region support cell proliferation exclusively in the permissive inner domain after 28 days. Construct size, 1.5x2 cm. G. Formation of 3D plant cell–based living materials. Representative photographs over 28 days of growth. Non-photosynthetic ELMs with coral (i), honeycomb (ii), and cubic (iv) architectures, while photosynthetic cells form honeycomb ELMs (iii), all printed using pec2alg*5 bioink over 28 days of growth. Coral structures base, 2 x 2 cm; honeycomb diameter of 6 cm (ii, iii); cubes, 1 x1 x 1 cm. H. Cell viability in honeycomb ELMs at day 28 of growth. Representative confocal Z-stack maximum intensity projections show live cells in cyan (fluorescein diacetate, FDA) and dead cells in red (propidium iodide, PI) for non-photosynthetic (top) and photosynthetic (bottom) ELMs. Scale bar, 2mm. I. TEM micrographs of plant cell wall-hydrogel interface from ELMs based on pec3.5alg2.5 and pec2alg*5 bioinks.

Rheological characterization revealed pronounced shear-thinning behaviour across all formulations, with viscosity decreasing with increasing shear rate, thereby facilitating extrusion printing. Among the formulations designed for 2D structures, pec3.5alg2.5 (wt/v%) was selected because it exhibited the best printability, characterized by strong shear-thinning behavior (n=0.45), continuous filament formation and excellent shape fidelity.

Formulations optimized for 3D printing, which contained high-viscosity alginate, exhibited even lower shear-thinning indices (n = 0.06–0.26), indicating a more pronounced decrease in viscosity with increasing shear rate (Fig. 2C, Fig. S3 and Supplementary Table 1). Both the storage (G′) and loss (G″) moduli of these formulations increased relative to the pec3.5alg2.5 ink, and they exhibited a distinct G′–G″ crossover point, indicative of improved structural recovery and shape retention following extrusion (Fig. 2D and Fig.S2, S3).The suitability of these formulations for centimetre-scale 3D printing was demonstrated by the fabrication of cm³ cubes with high shape fidelity (Fig. 2B, Fig. S2). Following initial assessment of cell viability in these structures, pec2alg*5 was selected as the optimal formulation, balancing centimetre-scale printability, shape fidelity and long-term cell viability (Fig.S4-S8).

We next investigated the ability of the bioprinted constructs to support sustained growth of non-photosynthetic and photosynthetic cells over several weeks and thereby form plant-based engineered living materials. Non-photosynthetic cells proliferated throughout 2D mesh and solid-square architectures while maintaining structural integrity and shape fidelity (Fig. 2E i, E ii, Fig.S9). Photosynthetic cells also exhibited robust growth within 2D mesh structures; however, their proliferation extended beyond the boundaries of the original printed geometry (Fig. 2E iii, Fig.S9).

Hydrogel composition also provided a means to control cell growth within printed constructs. We hypothesized that by doubling the concentrations of both pectin and alginate (pec7alg5) it will result in a denser hydrogel network that will prevent cell proliferation. Constructs were therefore printed with an inner region composed of pec3.5alg2.5 bioink and an outer region composed of abiotic pec7alg5 ink. Indeed, we observed selective proliferation only within the permissive inner matrix, enabling spatial control of cell growth and programmed pattern formation within a single living construct (Fig. 2F and Fig. S10).

The capacity of plant cells to form 3D ELMs with complex architectures was subsequently examined (Fig. 2G and Fig. S11). In ∼ 1 cm³ cubic constructs, viable cells were predominantly located at the area close to the surface, while only a few cells remained viable in the center (Fig. S7-S8). Similarly, coral-like structures despite having more open architecture than cubes, exhibited poor long-term viability, with cell growth largely confined to the surface of the constructs (Fig. S11-S12). In contrast, honeycomb ELMs with non-photosynthetic (Fig. 2G ii) and photosynthetic cells (Fig. 2G iii) with a diameter of 6 cm maintained high cell viability after 28 days, with FDA-stained living cells dominating and only minimal PI-stained dead cells observed throughout the structures (Fig. 2H, Fig. S13, S14, S15). These findings indicate that open architectures with reduced diffusion distances facilitate nutrient transport and support long-term cell viability and proliferation within 3D plant-based ELMs.

TEM analysis of both 2D and 3D ELMs revealed typical anatomical features of cells within the hydrogel matrices (Fig. S16). Examination of the interface between the plant cell wall and the surrounding hydrogel showed no discernible gaps, indicating intimate contact between the two components at the nanoscale (Fig. 2I, Fig S16). The hydrogel matrix appeared to seamlessly integrate with the cell wall, suggesting that the biomimetic environment supports close cell–matrix interactions and promotes incorporation of living cells into the material architecture.

These findings establish extrusion bioprinting as a robust platform for fabricating plant-based living materials with controllable architecture and growth.

### Plant cell wall biocatalysis enables in situ formation of electronically conductive living cells

To further demonstrate the potential of plant cells for engineered living materials, we introduced non-native electronic functionality into living cells. Previous studies have shown that endogenous plant peroxidases can catalyse the in vivo polymerization of conjugated trimers, enabling seamless integration of conducting polymers within plant tissues^30,31^. Following a similar strategy, we leveraged endogenous peroxidase activity to catalyse the polymerization of the conjugated trimer ETE-S into the conducting polymer p(ETE-S) directly on the cell wall (Fig. 3A). Toxicity assays confirmed that ETE-S concentrations up to 100 μM were well tolerated across different cell growth stages, with minimal cell death observed (Fig. 3B). Higher concentrations of ETE-S caused cell damage and death, as indicated by staining assay. (Fig.S17) Microscopy confirmed successful in-vivo polymer formation, with cells exhibiting variable thickness surface coatings after 24 h incubation, indicating heterogeneous functionalization efficiency (Fig. 3C).

**Figure 3:**
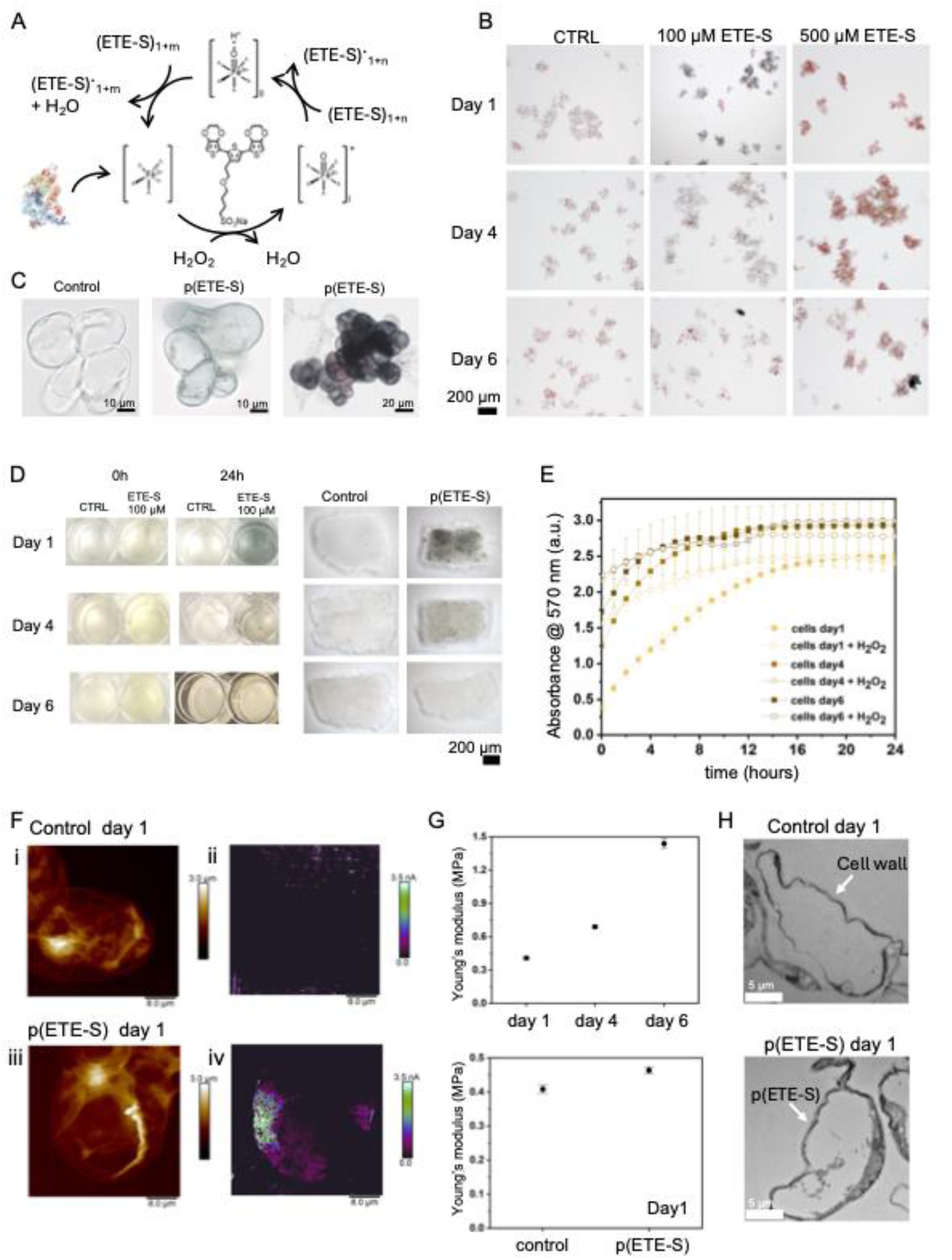
Harnessing plant cell biocatalytic machinery enables formation of living conductive cells. A. Schematic of peroxidase-catalysed radical polymerization of the ETE-S conjugated trimer, resulting in formation of the conductive polymer p(ETE-S). H_2_O_2_ oxidizes peroxidase to its active intermediate, compound I. Compound I subsequently oxidizes one ETE-S molecule and is reduced to compound II. The enzyme then returns to its resting state by oxidizing a second ETE-S molecule, concomitantly producing water. The resulting ETE-S radicals couple to form dimers. Further polymerization can proceed either through the entry of ETE-S oligomers into the peroxidase catalytic cycle or through radical transfer from ETE-S radicals to existing oligomers, generating new radical sites that undergo oxidative coupling to produce progressively larger oligomers (adapted from^29^). B. ETE-S toxicity assay in non-photosynthetic cells after 24h incubation at 100 and 500 μM for cells at different growth stage. 24h after incubation dead cells are stained with erythrosin B and appear pink. C. Micrographs of control and in vivo ETE-S-functionalized day 1 cells showing thinner (middle) and full (right) p(ETE-S) coating after 24 h incubation, indicating heterogeneous surface functionalization. Scale bars, 10 and 20 μm. D. Functionalization efficiency as a function of cell age (in respect to subculture day). Cells at day 1, 4, and 6 are shown before and after 24 h incubation with 100 μM ETE-S. Day 1 and day 4 cells exhibit stronger darkening, indicating enhanced polymer formation as shown in the well plates and in the stereomicrographs of cell aggregates before and after treatment. Scale bar, 200 μm. E. Endogenous peroxidase activity measured over 24h using the Amplex red assay for days 1, 4, and 6 cells with and without addition of 10 μM H_2_O_2_. Day 1 cells show lower activity, while exogenous H_2_O_2_ accelerates Amplex red conversion in all cases, indicating limited endogenous H_2_O_2_ availability. F. AFM topography (i, iii) and current maps (ii, iv) of PFA fixated control cells and p(ETE-S)-functionalized cells on day 1 (after 24h of incubation). Functionalized cells exhibit measurable current. G. Young’s modulus of control living cells at days 1, 4, and 6, showing increased stiffness with cell age. H. Young’s modulus of control and p(ETE-S)-functionalized living day 1 cells, showing a slight increase (11.8%) in stiffness upon functionalization. I. TEM images of control and p(ETE-S)-functionalized cells at day 1, showing integration of the p(ETE-S) layer with the cell wall. Scale bar, 5 μm.

Day 1 cells exhibited the darkest colour, indicative of the formation of a thicker polymer layer around the cell wall. The intensity of this dark coating decreased in day 4 cells, corresponding to a thinner polymer layer, whereas no obvious coating was observed in day 6 cells. Functionalization was therefore strongly dependent on cell age, with day 1 cells exhibiting the most effective polymer formation and day 6 cells the least (Fig. 3D). To determine whether this effect was related to peroxidase activity, endogenous peroxidase activity was assessed using the Amplex red assay in the presence of endogenous H₂O₂ only. Surprisingly, day 1 cells were less efficient at oxidizing the Amplex red reagent (Fig. 3E, Fig S18, Fig. S19), indicating a lower overall oxidizing capacity despite exhibiting more extensive ETE-S polymerization. This apparent discrepancy can be explained by differences in polymerization kinetics. Lower peroxidase activity may favour the formation of longer oligomers through sustained radical transfer, whereas higher peroxidase activity generates radicals more rapidly, promoting the formation of shorter oligomers and limiting extended polymer growth^29^. The addition of exogenous H₂O₂ increased the apparent peroxidase activity in all cases, indicating that endogenous H₂O₂ availability limits reaction kinetics within the cells (Fig. 3E, Fig S18, S19). Interestingly, this limitation may be beneficial for polymerization efficiency, as slower radical generation appears to favour the formation of longer conductive oligomers.

Conductive-AFM confirmed that functionalized cells acquired conductive properties. While control cells exhibited no measurable current, p(ETE-S) functionalized cells showed currents up to 3.5 nA, with spatial heterogeneity reflecting uneven polymer deposition or variable contact with the bottom electrode (Fig. 3F).

Mechanical measurements revealed a progressive increase in Young’s modulus with cell age by 240% at day 6 compared to day 1, consistent with cell wall stiffening during maturation^32–34^ (Fig. 3G, Fig S20, S21). Polymer functionalization induced a modest additional increase in stiffness (11.8%), suggesting reinforcement of the polysaccharide network without disrupting native architecture (Fig. 3G, Fig S20, S21). TEM further confirmed the presence of the conducting polymer close to the cell wall matrix (Fig. 3H, Fig. S22). To preserve this polymer coating during sample preparation, a milder embedding protocol was employed for cell sectioning, resulting in lower image quality. Collectively, these results demonstrate that plant cells can act as biofactories for the in-situ formation of conducting polymers, enabling living cells with coupled mechanical and electronic functionalities.

### Engineered living materials exhibit adaptive, evolvable and programmable properties

We next examined whether plant-based ELMs retain the ability to dynamically change their properties over time, forming 4D living materials that integrate adaptive, evolvable, and programmable behaviors (Fig. 4A). Non-photosynthetic ELMs transitioned to a photosynthetic state upon light exposure and reduced sucrose content, at day 14 of growth, demonstrating an environmentally triggered metabolic shift. Chlorophyll autofluorescence confirmed the formation of functional chloroplasts within the ELMs. This transition represents an adaptive response, as cells switch to a metabolically favorable state by developing chloroplasts that enable photosynthesis. At the same time, this transition can be externally controlled, illustrating programmable regulation of metabolic function.

**Figure 4:**
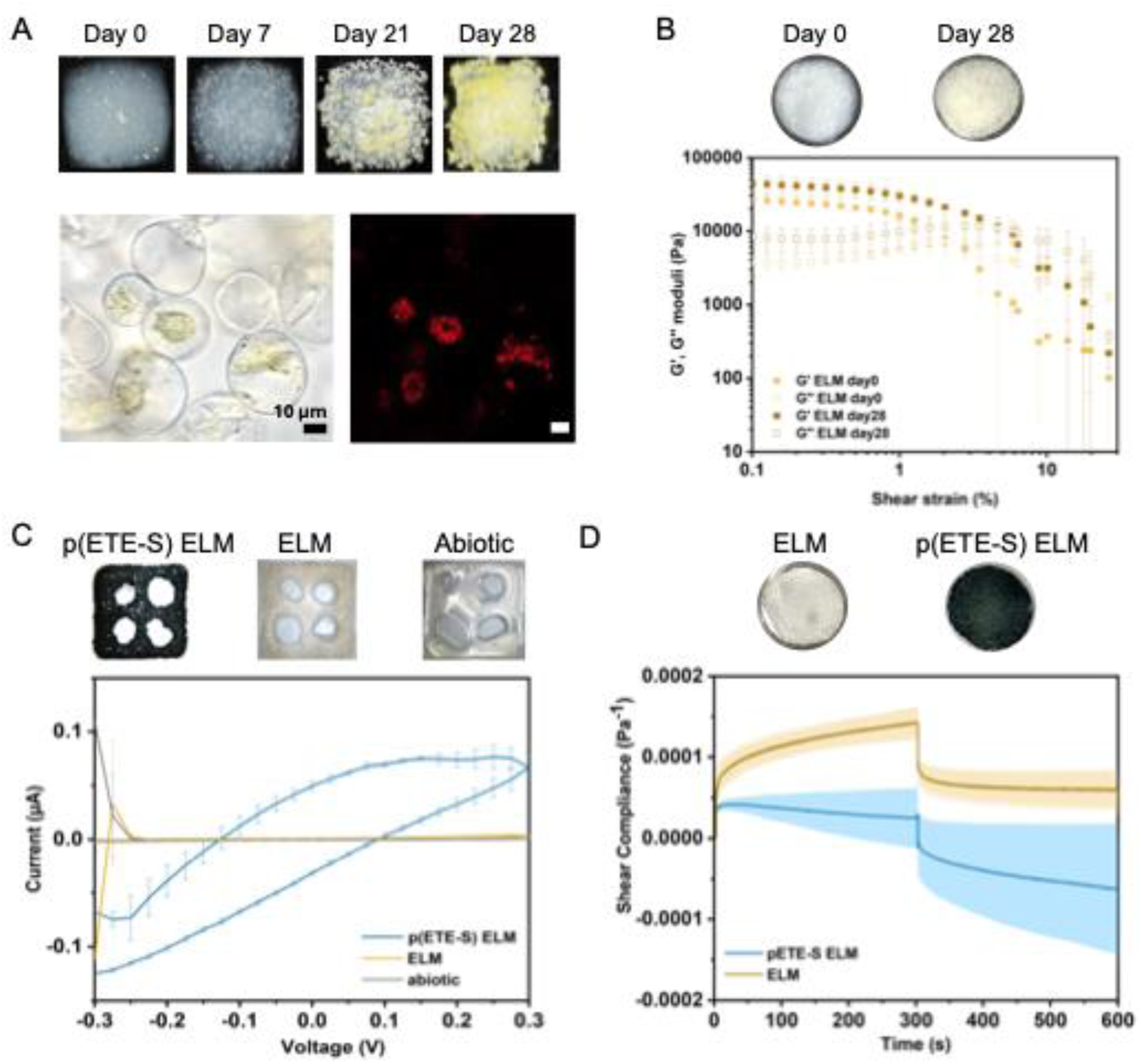
4D ELMs: plant-based living materials with adaptive, evolvable and programmable properties. A. Adaptive properties. Non-photosynthetic ELMs transition to a photosynthetic state upon light exposure at day 14, reflecting a metabolically beneficial adaptive response. Brightfield and confocal fluorescence micrographs show photosynthetic cells within the ELM. Chlorophyll autofluorescence is shown in red. ELM size, 1x1x cm. Scale bar, 10 μm. B. Evolvable rheological properties. Non photosynthetic ELMs based on pec3.5alg2.5 bioink (disc geometry; diameter, 2 cm; thickness, 2.0 ± 0.5 mm) show evolvable rheological properties. Storage (G′) and loss (G″) moduli as a function of shear strain are shown at day 0 and after 28 days. Cell proliferation drives changes in viscoelastic properties, resulting in increased moduli and shift in cross over point. C. Programmable electronic and electrochemical properties. ELMs functionalized with ETE-S via endogenous peroxidase-mediated polymerization exhibit macroscopic conductivity and charge storage, as demonstrated by two probe current-voltage characterization (−0.3 to +0.3 V, scan rate 50 mV s⁻¹). ELM size, 1x1 cm. D. Modulation of rheological properties via cell-mediated polymerization. Creep– recovery tests (10 Pa applied stress; 5 min creep, 5 min recovery) show that p(ETE-S)-functionalized ELMs exhibit more resistance to deformation and more plastic behavior compared to non-functionalized controls.

We next investigated whether cell growth alters the rheological properties of the ELMs. Rheological analysis revealed that cell presence and proliferation modify mechanical behavior over 28 days, indicative of evolvable properties driven by intrinsic biological activity (Fig. 4B). ELMs based on pec2.5alg3.5 after 28 days of growth exhibited increased storage (G’) and loss moduli (G’’) from day 0, indicating that cells presence acts as a reinforcement agent. The G’ increased by 65% (from 26.9 kPa at day 0 to 44.3 kPa at day 28) while the G” increased by more than 100% (from 3.8 kPa at day 0 to 8.5 kPa at day 28). The strengthening of the ELM over time reflects progressive network reinforcement mediated by cell presence, proliferation, and matrix remodeling. We hypothesize that cells form more junction zones between biopolymers and their cell walls, increasing entanglements, and strengthening the material. Furthermore, we observed the evolution of the G’–G” crossover point from 2.3% to 4.9% strain. The presence of cells brings more elastic properties and more resistance to deformation. When using the gel composition suitable for 3D structures, with the high viscosity alginate formulation pec2alg*5, we observe similar trends but smaller overall changes in the rheological properties (Fig. S23). This indicates that the evolving properties depend on cell-matrix interactions.

In addition, the biocatalytic activity of plant cells enabled programmable modulation of material properties. ELMs acquired electrochemical functionality through enzyme-mediated polymerization of conjugated trimer ETE-S and formation of p(ETE-S). Current– voltage measurements revealed the emergence of macroscopic electrical conductivity and charge-storage capacity, evidenced by the hysteresis observed in the functionalized ELMs, confirming the translation of cellular-scale biochemical processes into macroscopic electronic properties (Fig. 4C, Fig S24, Fig. S25).

Functionalization with p(ETE-S) also modified the mechanical properties of the material. Under an applied stress of 10 Pa, p(ETE-S)-functionalized ELMs exhibited lower compliance than unfunctionalized controls, indicating increased resistance to deformation (Fig. 4D). However, during the recovery phase, the p(ETE-S)-ELMs did not fully return to their initial state, demonstrating reduced elastic recovery and increased permanent deformation. Consistent with this observation, the crossover between the storage (G′) and loss (G″) moduli occurred at lower shear strains, indicating an earlier transition from solid-like to liquid-like behavior (Fig. S26). These results suggest that the conductive polymer coating formed at the cell surface enhances cell–matrix interactions, reinforcing the composite while simultaneously reducing network recoverability and promoting a more plastic-like mechanical response (Fig. 4D).

Together, these results demonstrate that plant-based ELMs combine adaptive response to environmental cues, programmable functionality via externally applied stimuli, and evolution of material properties through internal biological processes, enabling a new class of dynamic, multifunctional living materials.

### Living materials for energy dissipation and photosynthetic activity

Next, we investigated the interplay between cellular activity and structural geometry in determining mechanical performance, revealing a geometry-dependent adaptive behaviour (Fig. 5A, B, Fig S27). ELMs in hexagonal honeycomb and re-entrant auxetic architectures exhibited distinct deformation responses under tensile loading. In auxetic ELMs, the presence of cells increased both maximum force required for first strut failure (0.91 N vs 0.35 N) and extensibility (8.61 mm vs 6.73 mm) compared to abiotic controls (Fig. 5C, fig. S28). In contrast, honeycomb ELMs failed at lower forces (0.34 N vs 0.58 N) and extensibility (7.1 mm vs 15.1 mm) (Fig. 5C, fig. S29). Under tensile loading, the ribs in a honeycomb structure deform primarily through axial stretching and slight bending. The presence of the cells introduces heterogeneities and local stiffening leading to premature failure. In contrast in re-entrant auxetic structures the inward-facing ribs unfold under stretching through a deformation mechanism initially dominated by rotation of re-entrant hinges, followed by bending of inclined ribs. In this case, cells partially constrain rib bending and hinge rotation thereby acting as reinforcing elements that enhance the load-bearing capacity during deformation. These results demonstrate that the mechanical contribution of cells is not only intrinsic but emerges from their interaction with structural geometry, highlighting a dynamic coupling between biological activity and mechanical design.

**Figure 5:**
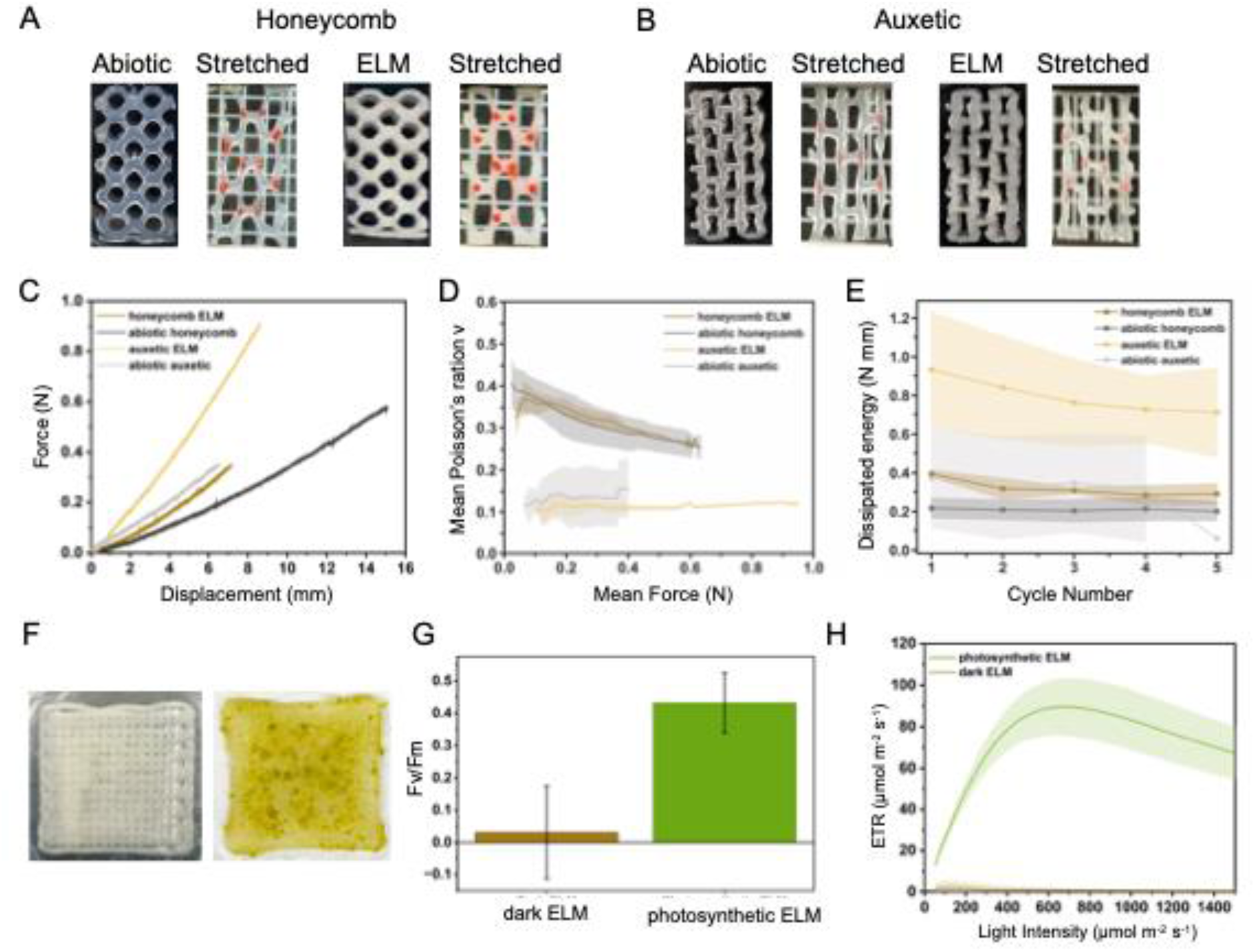
Living mechanical metamaterials and photosynthetic engineered living materials. A., B. Structural deformation of ELMs under tensile loading in (A) honeycomb and (B) auxetic architectures. Panels show the corresponding abiotic structures, ELMs after 14 days of growth, and their response under applied tensile stress. C. Force-displacement curves of honeycomb and auxetic ELMs under tensile loading to failure (average value, n = 3; sample size, 4.5 x 2.5 cm). Auxetic ELMs exhibit higher failure force and greater extensibility than their abiotic counterparts, whereas honeycomb ELMs fail at smaller forces and extensibility compared to abiotic controls. D. Poisson’s ratio as a function of applied force for auxetic and honeycomb ELMs and their abiotic counterparts. Honeycomb structures exhibit higher positive Poisson’s ratios that decrease with increasing force, whereas auxetic structures maintain low and relatively constant values over a force range. E. Energy dissipation as a function of tensile cycle. Auxetic ELMs display higher and more stable energy dissipation than honeycomb structures. F. Photograph of non-photosynthetic (dark) and photosynthetic ELM, sample size 4x4cm at 21 days. G. Maximum quantum yield of photosystem II, Fv/Fm of photosynthetic ELMs and non-photosynthetic ELMs as control. Photosynthetic ELMs exhibit measurable photosynthetic activity. H. Fitted electron transport rate curves of photosynthetic ELMs and non-photosynthetic ELMs controls as a function of actinic light intensity. Photosynthetic ELMS show characteristic light-dependent increase in ETR.

Poisson’s ratio measurements further confirmed these differences, with auxetic ELMs maintaining a stable near-zero Poisson’s ratio over a broad force range, whereas honeycomb structures exhibited decreasing positive values with increasing load (Fig. 5D, Fig. S30). Although ideal auxetic structures exhibit negative Poisson’s ratios, this behavior is highly dependent on geometric fidelity, including the shape of the unit cells and the degree of corner rounding. Simulations have shown that increasingly rounded unit cells can shift the Poisson’s ratio towards non-negative values^35^. The near-zero values observed here indicate that the material can undergo longitudinal stretching with minimal transverse deformation and contraction.

Energy dissipation analysis revealed that auxetic ELMs sustain higher and more stable dissipation across loading cycles, indicating enhanced energy absorption and fatigue resistance (Fig. 5E, S31, S32). In honeycomb structures, instead, the influence of cells was less pronounced. The incorporation of cells increased the dissipated energy compared to abiotic honeycomb hydrogels; however, this increase was moderate and did not reach the values observed in auxetic ELMs. The enhanced energy dissipation of auxetic ELMs likely arises from the coupling of the intrinsic deformation mechanisms of the auxetic architecture with additional dissipative mechanisms introduced by the cells, including dynamic cell–matrix adhesion, intrinsic cellular viscoelasticity, and cell-mediated load transfer.

Overall, these results demonstrate that the cells presence enhances energy dissipation in hydrogels, but the extent of this effect is strongly influenced by the underlying architecture, with auxetic structures showing the most pronounced cell-mediated increase.

Finally, we evaluated photosynthetic activity of centimeter-scale ELMs (Fig. 5F, Fig. S33). The maximum quantum yield of photosystem II (Fv/Fm) reached values of 0.43 ± 0.09, indicating the presence of functional photosynthetic machinery within the ELM, whereas non-photosynthetic ELMs exhibited negligible values, consistent with the absence of photosynthetic activity (Fig. 5G). Negative Fv/Fm values arise because Fo and Fm are nearly identical and approach the detection limit of the instrument, where background fluorescence and light-scattering contributions become significant. Photosynthetic performance was further quantified using rapid light curves. Photosynthetic ELMs displayed a characteristic light-dependent increase in electron transport rate (ETR), reaching a maximum value of 89.3 ± 13.5 µmol m⁻² s⁻¹ at an irradiance of 655 µmol m⁻² s⁻¹ (Fig. 5H). At higher light intensity, ETR decreases potentially due to deactivation of non-photochemical quenching. In contrast, non-photosynthetic ELMs exhibited ETR values close to zero and no light-dependent response across the entire irradiance range. Together, these results demonstrate that plant cells retain photosynthetic ability throughout ELM formation and confer this functionality to the macroscopic material, enabling efficient light-driven electron transport within centimeter-scale living constructs.

## Discussion

Engineered living materials seek to bridge the gap between static engineered matter and the dynamic behavior of biological systems. Here, we establish plant cells as a distinct class of building blocks for engineered living materials and demonstrate that their intrinsic metabolic plasticity, cell-wall biochemistry and mechanics can be harnessed to create materials that exhibit adaptive, evolvable and programmable properties. By integrating plant cell culture, bioprinting, organic electronic materials and mechanical metamaterials within a single platform, our work expands the design space of photosynthetic ELMs beyond current microorganism and organelle-based approaches. A defining feature of biological organisms is their ability to continuously adapt metabolism to environmental conditions. We show that non-photosynthetic wild type *Arabidopsis thaliana* cells embedded within macroscopic materials can transition to a photosynthetic state in response to light while maintaining viability and biomass accumulation. Unlike most existing ELMs, where function is predetermined by the chosen organism or engineered circuitry, plant cells exploit their metabolic plasticity to acquire new functionality in response to environmental cues. This behavior more closely resembles adaptation in natural tissues and suggests a route towards materials whose function is not fixed at fabrication but can emerge during deployment.

The inherent mechanical properties of plant cells due to the cell wall and their ability to interact with their local environment further enabled the emergence of evolvable material properties. Continuous cell growth and proliferation altered the rheological properties of the ELM over time, resulting in progressive reinforcement of the hydrogel network. Previous photosynthetic ELMs have primarily achieved mechanical reinforcement through the formation of secondary structural phases, including photosynthesis-driven matrix crosslinking^19,20^, biomineralization^17,36^, or symbiotic growth with cellulose producing bacteria^16^. By contrast, the mechanical evolution observed in our plant-based ELMs arises without mineral deposition or synthesis of a new structural extracellular component. Instead, growing plant cells progressively become an active structural element of the composite, with cell proliferation, cell-wall maturation and cell–matrix interactions driving changes in viscoelastic properties over time.

Beyond exploiting native biological processes, we demonstrate that plant cells can serve as biofactories for the generation of non-native functionality. Endogenous cell-wall peroxidases catalyzed the in-situ formation of the conducting polymer p(ETE-S), producing electronically active living cells without genetic modification. Previous studies have shown the integration of conducting polymers within intact plants^30,31,37^; however, their incorporation into living materials has remained unexplored. The resulting macroscopic ionic–electronic conductor demonstrates how cell wall machinery can be coupled with synthetic functional materials to create programmable behavior. Importantly, polymer formation occurred within the cell-wall environment while largely preserving cellular viability and mechanical integrity. This approach establishes a strategy for translating cellular biochemical activity into macroscopic electronic function and suggests opportunities for future living bioelectronic interfaces, sensing systems and adaptive energy materials.

The multifunctionality demonstrated here emerges from three distinct but complementary mechanisms: adaptation through environmentally triggered metabolic reprogramming, evolution through continuous biological growth and matrix remodeling, and programmability through externally initiated biofabrication processes. Although these concepts are often discussed interchangeably within the ELM field, our results highlight important distinctions. Adaptation enables materials to respond autonomously to environmental conditions, evolution reflects gradual changes arising from intrinsic biological activity, and programmability allows the deliberate introduction of new functionality through external intervention. Plant cells uniquely combine all three capabilities within a single living platform, providing a conceptual framework for designing future generations of 4D living materials.

Our results further reveal that the functionality of living materials is governed not only by cellular activity but also by architectural design. The contrasting behavior observed between auxetic and honeycomb ELMs demonstrates that geometry determines how biological reinforcement is translated into macroscopic mechanical performance. In auxetic structures, cells enhanced both strength and extensibility while increasing energy dissipation and maintaining a stable Poisson’s ratio. In contrast, the same biological component reduced mechanical performance in honeycomb architectures. These observations suggest that living cells should not simply be regarded as active fillers within a passive scaffold. Rather, their contribution depends strongly on how biological processes interact with local stress distributions and deformation mechanisms. The concept of geometry-dependent biological reinforcement introduces a new design principle for living metamaterials, where function emerges from the coupling between architecture and cellular activity.

Beyond their evolving mechanical behaviour, plant-based ELMs offer a route towards structurally integrated photosynthetic materials that combine biological energy conversion with macroscopic function. A central challenge for such systems is maintaining photosynthetic performance while forming an engineered structure. The maximum quantum yield of photosystem II (Fv/Fm) in our photosynthetic ELMs was lower than that of wild-type *Arabidopsis thaliana* leaves (≈ 0.8) but remained within the range commonly reported for hydrogel-encapsulated photosynthetic microorganisms^38–41^. Reduced Fv/Fm values likely arise from a combination of self-shading, light scattering, and prolonged culture within the hydrogel environment^41–44^. Despite this reduction, the maximum electron transport rate reached 89.3 ± 13.5 µmol m⁻² s⁻¹ is comparable to values reported for wild-type *Arabidopsis thaliana* and substantially higher than those typically observed in algae- and cyanobacteria-based photosynthetic living materials^39,40,44^. Interestingly, maximum ETR was reached at a higher irradiance (655 µmol m⁻² s⁻¹) than in native leaves (≈ 300 µmol m⁻² s⁻¹), consistent with light attenuation within the hydrogel matrix. The ability to sustain photosynthetic activity at levels approaching those of native plant tissues, despite the absence of leaf-level organization and vascular transport networks, highlights the robustness of plant cells as photosynthetic building blocks. More broadly, these findings suggest that plant-based ELMs can translate cellular photosynthetic function to the material scale, opening opportunities for autonomous living materials capable of oxygen production, carbon capture and biologically powered functionalities.

Several limitations remain to be addressed. Long-term maintenance of photosynthetic performance will require optimization of light transport, nutrient delivery, and construct architecture, particularly in larger three-dimensional structures. Furthermore, while the present study demonstrates adaptation, evolution, and programmability at a laboratory scale, future work should investigate how these materials perform in open and dynamic environments. Integrating plant-based ELMs with synthetic biology tools, developmental control strategies and autonomous feedback mechanisms could further enable materials capable of more sophisticated decision-making and environmental interaction.

In conclusion, our work establishes plant cells as versatile and previously underexplored chassis for engineered living materials. By combining metabolic plasticity, cell-wall biocatalysis, cell-matrix interaction, photosynthetic and structural function, we demonstrate living materials that adapt, evolve, and acquire new functionality over time. These findings expand the scope of photosynthetic engineered living materials beyond current microbial and organelle-based systems and lay the foundation for a new generation of plant-based living matter.

## Methods

### Materials

The following chemicals were purchased and used as received: pectin from citrus peel (TCI), sodium alginate (Sigma Aldrich), alginic acid sodium salt (high viscosity; ThermoScientific), agar (microbiology tested; Sigma Aldrich), Murashige & Skoog (MS) medium (including vitamins; DuchefaBiochemie), calcium chloride dihydrate ≥99% (Sigma Aldrich), poly-L-lysine solution (0.1% w/v in H_2_O; Sigma Aldrich), formaldehyde solution 4% (buffered, pH 8.3; Sigma Aldrich), glutaraldehyde solution (25% stock, EM grade; Taab Laboratories Equipment Ltd, G011/2), phosphate (PB) buffer (Supelco), hydrogen peroxide solution (30wt% in water, contains inhibitor, ACS reagent. Sigma Aldrich), sucrose (≥99.5%; Sigma Aldrich), fluorescein diacetate (Sigma Aldrich), propidium iodide (ThermoFisher), erythrosin B (Sigma Aldrich), Amplex red hydrogen peroxide/peroxidase Assay kit (ThermoScientific), bis[3,4-ethylenedioxythiophene]-3-thiophene butyric acid, sodium salt, trimer (ETE-S) was synthesized by Dr. Daniele Mantione at POLYMAT, University of the Basque Country

### Preparation of plant cell cultures

*Arabidopsis thaliana* cell cultures were generated from a solid back up agar plate provided by Prof. Åsa Strand, Umeå Plant Science Center. Under an operating biosafety cabinet, dark cells were scraped off the agar plate using a spatula and transferred in each well of a 6-well plate containing 5 mL of MS medium 3% sucrose. After a gentle homogenization with a pipette, the plate was sealed with micropore tape and put on the orbital shaker at 120rpm and 22°C in the incubator in dark conditions. The plate was checked every 3 days until the density was high enough. After approximately 10 days, 3 mL of cells were transferred with a sterile pipette in a 100-mL Erlenmeyer flask with 27mL of fresh MS medium 3% sucrose and placed on the same shaker in the incubator. Every 7 days, the cells were subcultured in a new flask in the ratio 1:10.

For generating photosynthetic cell cultures, 3 mL of dark cells were transferred in a new flask containing 27 mL of MS medium 1% sucrose and placed in light conditions (light intensity 150 μmol m^2^ s, white 4000K LED panel) to induce the development of chloroplasts, according to the established protocol^25^. Every 7 days, first the medium was replaced with fresh MS 1% sucrose by placing the flask on a 60° tilted 3D printed support, and then the photosynthetic cells were transferred in a new flask with the ratio 1:10. 250-mL Erlenmeyer flasks containing 72 mL of MS medium with either 3 or 1% sucrose and 8 mL of non-photosynthetic (dark) or photosynthetic cells respectively were used for the growth curves.

### Growth curves

Two methods were used to characterize the growth of plant cell cultures, cell wall sedimentation (CVS) and dry weight (DW), according to the protocol reported In ref ^45^.

*Cell wall sedimentation method*: the measurement has been performed for 3 biological replicates of cell cultures with 3 technical replicates each, by using the 3D printed 60° tilted support with an imprinted ruler. Every day for 14 consecutive days the 250-mL Erlenmeyer flasks containing the cell cultures were placed on the support for 5 minutes and the height of the cells’ sediment was registered. The height was then converted to volume by using the corresponding volume of water which then represented the CVS value. The CVS value was then converted for 1L of suspension and plotted versus time to obtain the CVS growth curve.

*Dry weight (DW) method*: paper filter discs with a porosity of 100 μm (VWR) were previously dried in oven at 60°C to remove any residual humidity. By using a Büchner vacuum funnel, 1 mL of cells was collected by each flask and filtered on a pre weighed paper filter disc, then placed in oven at 60°C for 24 hours. The filter with dried cells was weighed, registering the stable weight, and subtracted the weight of the filter paper to obtain the DW of the cells. The ΔDW weight was then plotted versus time to obtain the DW growth curve.

The two methods were performed on dark, photosynthetic and transitioned from dark to light cell cultures.

### Preparation of bioinks and 3D Bioprinting

The 3D Bioprinting process was performed using Cellink (BioX Gen3) and the software DNA Studio 4 for the design of the patterns. Sterile 3mL cartridges with pistons, female-female luer lock adapters and different diameter size nozzles were purchased from Cellink.

Inks with different concentrations of biopolymers (wt/v%) were prepared by weighing the powders and dissolving them in autoclaved (121°C) MS medium with 1 and 3 wt/v% sucrose concentrations for photosynthetic and dark cells, respectively. The solutions were kept under UV light for 1 hour and then under stirring at 400 rpm at 40°C overnight. Prior to the printing process, glass petri dishes, nozzles and cartridges, washing and storage solutions, were sterilized under UV light for one hour.

Prior to printing, up to 4 mL of cells 2 days after subculturing (denoted day 2 cells) were transferred in a sterile beaker and the excess of MS medium was removed. Then, 3mL of ink was added and gently integrate the cells with a spatula. Once homogenized, the bioink was transferred in the cartridge and it was ready for bioprinting. After the printing process in a glass petri dish, the abiotic and the ELM hydrogels were immersed for 2 to 4 minutes, depending on the size, in autoclaved 100 mM CaCl_2_ as crosslinking agent, then washed three times with autoclaved MS solution, then transferred in vials or petri dishes with MS medium and placed in light (for photosynthetic cells) or dark (for non-photosynthetic cells) incubators at 22°C for growth until further characterization.

### Atomic Force Microscopy (AFM)

The glass slides were cleaned with 70% ethanol solution prior any treatment. In the biosafety cabinet a solution of poly-L-lysine (PLL) 0.1 w/v% was filtered directly on the glass slides by using syringe filters with mesh size of 20 μm and left in closed petri dishes for 20 minutes. After this time, the excess of the solution was removed and the glass slides left to dry overnight.

The samples for the mechanical AFM were prepared by casting on the PLL-coated glass slides the living dark cells at different growth stages (1, 4, 6 days after subculturing). After 24 hours, the glass slides were washed gently twice with fresh MS medium 3% sucrose to remove unattached cells.

For the C-AFM, first the glass slides underwent a UV-ozone treatment for 35 minutes. Gold electrodes for C-AFM were prepared by thermal evaporation of 5 nm Cr and 35 nm Au on each glass slide.

The gold coated glass slides were then prepared with poly-L-lysine as described above. The following day, living dark cells 1 day after subculturing were casted on the glass slides for 24 hours. After this step, the glass slides were washed with fresh MS medium 3% sucrose twice and then incubated for 15 minutes with 5 mL of PFA 4% solution in the fume hood. Then the slides were washed with fresh MS medium and then gently dried with N_2_ flux prior the analysis.

Conductive atomic force microscopy (C-AFM) was performed in a Dimension Icon XR microscope (Bruker). Height images and current maps were collected in PeakForce Tapping mode with Pt/Ir-coated silicon probes having a spring constant of 3 N/m, a resonance frequency of 75 kHz, and a nominal tip radius of 25 nm. For current maps a dc bias was applied to the conductive substrate while keeping the probe at ground.

AFM mechanical testing was performed by using calibrated probes having tip length 17 µm, tip radius 70 nm, and spring constant 0.1 N/m (PFQNM-LC-V2, Bruker). Measurements were performed by bringing the AFM probe in contact with individual cells at a controlled load force of 10 nN. 300 force curves were recorded per cell in an area of around 3 µm^2^, and around 10 cells per condition were measured. The elastic modulus was obtained by fitting the force curves through the Hertz model:

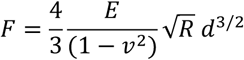

where F is the force, d is the indentation depth, E is the Young’s modulus, ν is the Poisson’s ratio, and R is the radius of the indenting probe. A Poisson’s ratio of 0.3 was used. All fittings were performed with the Nanoscope Analysis 2.0 software.

### Confocal Microscopy Imaging

2mL of living plant cells were stained with 10 μL of fluorescein diacetate (FDA, 5 mg/mL in acetone stock solution) and 10 μL of propidium iodide (PI, 1 mg/mL in water). After 10 minutes of incubation in dark conditions, 100 μL cells were casted on a glass slide and analysed in brightfield and fluorescence modes at the confocal microscope Zeiss LSM 980 with Airyscan 2. The samples were analysed in 5x, 20x and 63x (oil immersion) objectives with the following lasers: calcofluor (exc 405 nm, em 410-481 nm), fluorescein diacetate (exc 488 nm, em 490-550 nm), propidium iodide (exc 561 nm, em 561-650 nm), chlorophyll a (exc 639 nm, em 667-694 nm), erythrosin B (exc 525 nm, em 545-550 nm). Z-stack and tile scans were performed and then images were processed in ImageJ.

For the ELMs in cube shape, thin slices ware sectioned from the centre of the cube using a sterile scalpel at specific days after printing and each slice was immersed in a fluorodish with 2 mL of sterile MS medium 3% sucrose and stained as previously described for the cells.

For the ELMs in honeycomb design, an internal hexagon was cut out with a sterile scalpel. Each hexagon at a specific day after printing was immersed in a fluorodish with 2 mL of sterile MS medium 1/3% sucrose and then stained as previously described for the cells. The living cells with and without pETE-S were stained with 10 μL of erythrosin B 0.2 wt/v% in water and visualized in brightfield mode.

### Transmission electron microscopy (TEM)

Living cells 1, 4 and 6 days after subculturing were placed in sterile Eppendorf tubes with 2,5% glutaraldehyde (GA) in 0.1 M phosphate buffer (PB) pH 7.4, 4°C overnight. The solution was then replaced with 12% gelatine (Dr. Oetker) in 0.1M PB, pH 7.4, incubated at 37°C for 30 minutes. Each sample was then transferred to a fresh 12% gelatine solution and incubated for additional 10 minutes at 37°C. The samples were then placed on ice. The samples were sectioned, and transferred to 2.3 M sucrose (VWR 27480.294) overnight at 4 °C. Next day they were mounted to metal pins facing the surface with an extra drop of 2.3 M sucrose, to prevent ice crystal formation upon freezing, and instantly frozen in liquid nitrogen. To prepare cryosections, the samples were trimmed, and the presence of the cells were confirmed under a light microscope. Ultra-thin sectioning of approximately 70 nm thickness was done using an Ultra microtome (Leica EM UC7) and the sections were placed on hexagonal copper grids, with a formvar film and the carbon layer of approximately 2 nm. The grids with sections were incubated for 20 minutes at 37°C in 1X PBS (PBS Tablets Medicago 09-8912--100) pH 7,4 buffer to remove the gelatin. After that, the grids were washed with PBS five times for 2 minutes, followed by six consecutive washes in milliQ water for 1 minute. Contrasting was done by incubating the grids in methylcellulose (Sigma M-7140)/uranyl acetate (EMS England 22451), pH 4, on ice for 6 minutes. The grids were looped out using homemade Remanium wire loops and imaged on the transmission electron microscope (ThermoScientific Talos L120C TEM).

The alg2.5pec3.5 and pec2alg*5 ELMs 14 days after printing were fixed in 2.5% glutaraldehyde in phosphate buffer (NaH_2_PO_4_•H_2_O and Na_2_HPO_4_•2H_2_O), after post-fixed for 2h in 1% OsO_4_ in water. After that the samples were dehydrated in a graded ethanol series followed by resin infiltration (3:1,1:1,1:3 resin: pure ethanol) and finally embedded in pure Spurr’s resin (TAAB Laboratories Equipment Ltd, England). Ultra-thin sections of 70 nm thickness were stained with uranyl acetate and lead citrate and examined with the Thermo Scientific Talos L120C transmission electron microscope.

### Rheological characterization

The Malvern Kinexus rheometer with the rSpace software was used for the rheological analysis of inks and crosslinked abiotic samples and ELMs. A Peltier stainless steel bottom plate and a 20mm totally flat stainless steel top plate at 25°C were used for all the samples.

The shear thinning properties were investigated by measuring a shear rate table between 0.1-100 s^-1^, 10 samples per decade at 25°C and a gap of 0.8mm for all ink formulations. The amplitude sweep tests were run at strain between 0.1-1000% at 1 Hz, 25°C to investigate the viscoelastic properties of inks and at strain 0.1-100% with a gap of 2 ± 0.5 mm for the crosslinked abiotic and ELMs of disc shape (⌀ = 2 cm) at 0 and 28 days after printing.

The creep recovery tests were run for a creep phase by applying a stress of 10 Pa for 5 minutes, followed by the recovery phase for other 5 minutes.

The test was performed on duplicates, and the data were further processed on Origin 8.0.

### Electrochemical characterization

Linear sweep voltammetry with initial voltage of -0.3V and final voltage of + 0.3V, with a scan rate of 10, 25 and 50 mV/s was recorded three times for each of the 3 selected points for abiotic hydrogels, uncoated and pETE-S coated ELMs in dry and wet conditions at room temperature. SE-TG gold plated probe tips with diameter of 10 μm were purchased by Amerikanska Teleprodukter AB. With the use of micromanipulators, the tips were connected to the working and reference electrode cell cable terminations of the Gamry potentiostat Interface 1010E.

### Tensile stretching measurements

For the stretching test of auxetic and honeycombs abiotic and ELMs, a Thumbler GmbH tensile tester Z3 with a load cell model TCTN9110-3 kN was used.

Abiotic and ELMs with the two different architectures were printed in triplicates with sizes 4.5x2.5 cm, then incubated in MS medium 3% for 14 days in sterile sealed glass petri dishes in the incubators.

During the tensile test, time lapse images of samples’ deformation with load were captured using a high-definition Sony α6400 camera equipped with a Sony E 18-135mm f/3.5-5.6 OSS lens. High resolution images of 24 megapixels were acquired every 10 s during the stretching experiment. The images for each test were collected and converted in time lapse tagged image file format time lapse.

A custom Python pipeline based on OpenCV library was used to automatically detect red markers embedded in the hydrogel across time lapse frames, through color-based segmentation in HSV space, followed by contour detection to extract dots centroid positions. A frame-to-frame tracking based on nearest-neighbor matching was used to link consecutive markers over time, allowing the reconstruction of the dots’ trajectories over the tensile test recording. Marker trajectories and position coordinates were converted into physical units through spatial calibration and used for longitudinal and transverse strains calculation.

For each experiment, horizontal and vertical distances were measured between predefined marker pairs in each recorded frame (L(t)).

From these distances, engineering strain was calculated with respect to the first frame as the reference starting distance (gauge length L_0_), by using the generic formula:

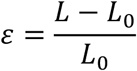

Transverse and longitudinal engineering strains, *ε_x_* and *ε_y_* respectively, were calculated between marker pairs and for each frame, as follows:

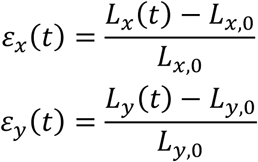

Poisson’s ratio was then calculated for each marker pair and averaged:

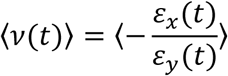

Data from different experiments were aligned to the same start time, interpolated onto a shared 10 s time grid while preserving the full duration of the longest experiment and summarized as mean ± standard deviation. Moreover, the first three aligned time points were excluded in trimmed plots to reduce the influence of initial tracking and loading artifacts.

Force data collected with the tensile tool were combined with the calculated Poisson’s ratio results to compare the deformation behaviour under loading. For each experimental group, force-time data were sorted and cleaned. A common overlapping time range for force measurement was defined between experiments, and force data were interpolated onto a common 10 s time grid. The first three time points were removed to match the trimmed Poisson analysis. Mean force and standard deviation were calculated across repeated measurements at each time point, after which the corresponding Poisson’s ratio and its standard deviation were interpolated onto the same time grid, allowing to direct match Poisson’s ratio with measured force.

For each specimen, the force-displacement curve was extracted and two characteristic maximum forces for the mechanical performance quantification were identified (*F_max_*). The first damage point was defined as the first significant force drop following a local maximum in the force-displacement curve. Instead, the maximum force point was detected as the highest force recorded during the traction test.

Hysteresis behaviour of the structures was evaluated by applying cycles of loading and unloading of the hydrogels, by directly imposing the maximum displacement and by measuring the corresponding force. Energy dissipation during the loading cycles was quantified from the force-displacement data. Both data sets were smoothed using a centered rolling median filter over a 2 s window to reduce experimental noise and prevent false cycle detection from signal fluctuations. The cycles were detected by peak detection from local maxima in the smoothed displacement dataset. Each cycle was defined as the interval between two consecutive displacement peaks, defined as *p_i_*, *p_i_*_+1_, with i as the cycle number. Following this, the dissipated energy for each cycle was calculated from the smoothed force-displacement curve in the form of absolute mechanical work, defined as the absolute area under the force-displacement curve:

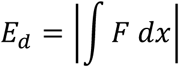

Numerically, the integral was computed using the trapezoidal rule, starting from force and displacement discrete data at each point j (*F_s_* (*j*) and *x_s_*(*j*)):

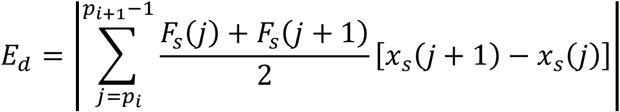

where *E_d_* is the dissipated energy for cycle, *F_s_* (*j*) and *F_s_*(*j* + 1) are the smoothed forces at two consecutive points, *x_s_*(*j*) and *x_s_*(*j* + 1) are the smoothed corresponding displacements, *p_i_* and *p_i_*_+1_ are consecutive displacement peak indices. Additionally, the absolute value was used to that energy loss was reported as a positive quantity. Energy values were grouped by cycle number for each experimental condition and the mean ± standard deviation was calculated across repeated samples.

All the previous measurements were repeated in independent experiments for auxetic and honeycomb structures, with and without cells.

### Stereomicroscope imaging

A Nikon SMZ1500 stereomicroscope was used to image cells with and without the pETE-S, abiotic and ELMs with and without pETE-S for the macro electrical characterization.

### Absorption spectroscopy

Absorption spectra were recorded by using the spectrophotometer PerkinElmer UV/Vis/NIR Lambda 1050+ coupled with an integrating sphere. Living photosynthetic plant cells at day 0, 7, 14, 21 and 28 days after subculturing were casted on glass slides by removing the excess of MS medium and directly analyzed. The spectra were recorded between 300 and 800nm, with interval of 4nm.

### Kinetics study

UV–Vis spectra were obtained using a microplate reader Biotek, Synergy H1 between 300 and 900 nm with a resolution step of 5nm at 23°C.

1 mL of living cells 1, 4 and 6 days after subculturing were incubated for 24 hours in dark conditions and under shaking with 100 μM of the trimer ETE-S and with and without 10 μM of H_2_O_2_ with a final volume of 2 mL in each well of a 12 well-plate.

For the Amplex red assay, the spectra were recorded 30 minutes after incubation and every hour for 24 hours, by setting an orbital shaking period of 5 seconds prior each recording.

Obtained data for triplicates were normalized to the minimum value before removing the MS medium as blank from the sample data.

### In vivo ETE-S polymerization and toxicity assay

1mL of living cells 1, 4 and 6 days after subculturing were incubated with specific volumes of the trimer ETE-S 2mM, previously autoclaved, to reach the final concentrations of 0, 10, 50, 100, 250 and 500 μM by keeping a final volume of 2 mL in each well of a 12 well-plate. After an incubation time of 24 hours in dark conditions and under shaking 1 mL of cells were stained with 10 μL of erythrosin B 0.2 wt/v% for 15 minutes in dark and then analyzed in brightfield mode at the confocal microscope. The experiment included triplicates for each concentration of the trimer.

### Amplex red assay

The Amplex red (AR) assay kit was purchased by Sigma Aldrich, and the solutions were prepared by following the included protocol. The working solution of 200 μM of Amplex red was prepared by diluting 100 μL of the 10 mM stock solution in 4.90 mL of the phosphate buffer 0.25M at pH 7.4. Then, a solution of 200 μM of AR reagent and 200 μM of H_2_O_2_ in 5 mL of buffer was prepared in a sterile glass vial by adding 100 μL of the 10 mM AR stock solution and 50 μL of the stock solution of H_2_O_2_ 20 mM. 100 μL of cells 1, 4 and 6 days after subculturing were added and a fixed total volume of 200 μL was kept in each well of a 96-well plate. The controls of the single reagents in the buffer solution, as well as of cells with AR 200 μM and H_2_O_2_ 200 μM were included in triplicates and incubated for 24 hours shaking in dark conditions at 22°C.

### Synthesis of ETE-S trimer

A detailed description of the synthesis and characterization of the ETE-S trimer was published elsewhere^46^.

### Photosynthetic characterization

The fluorometer integrated into the LI-6800 Portable Photosynthesis System (LI-COR Biosciences), equipped with the aquatic chamber (6800-18), was used to evaluate the photosynthetic performance of the engineered living materials. The chamber temperature was maintained at 22 °C, and the ELMs were dark-adapted for 30 min prior to the measurements. Samples (n=3 each condition) were cut in squares 2x2cm and placed inside the aquatic chamber using a plug accessory of the aquatic chamber. The measuring light was configured with a spectral composition of 90% red and 10% blue, while the actinic light intensity was initially set to 0 µmol m⁻² s⁻¹ (dark condition). A saturating pulse with a rectangular waveform, 1 s duration, 1 kHz data acquisition frequency, and an intensity of 6000 µmol m⁻² s⁻¹ was applied. Under dark-adapted conditions, the minimum fluorescence (F₀) and maximum fluorescence (Fₘ) values were recorded.

To determine the photosynthetic response to irradiance, actinic light was sequentially applied at 0, 50, 125, 150, 175, 300, 600, 900, 1200, and 1500 µmol m⁻² s⁻¹. At each irradiance level, fluorescence parameters were measured and the Electron Transport Rate (ETR) was calculated by the instrument software. For subsequent analysis, data were exported from the Excel files generated by the LI-6800 after each experiment. ETR values were calculated using the chamber cross section area (7.89 × 10⁻⁴ m²).

The maximum quantum yield of photosystem II (Fᵥ/Fₘ) was calculated according to:

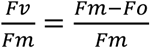

The model of Eilers–Peeters was used to fit the ETR curves^47^. Equation shows the form of the model where E is the light intensity.

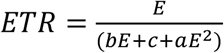

Curve fitting was performed in Python using the curve_fit function from the SciPy optimization package. From the fitted curve, the following photosynthetic parameters were derived:

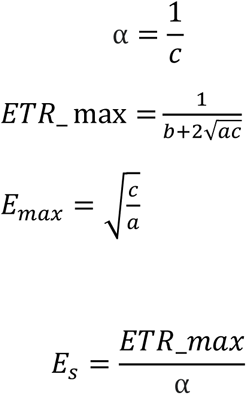

where, α is the initial slope of the ETR curve, ETR_max is the maximum value of ETR predicted by the model, E_max is the irradiance associated to ETR_max and E_s is the saturation irradiance.

Three independent ELM samples were analysed for each condition: abiotic, loaded with non-photosynthetic cells, and loaded with photosynthetic cells.

## Acknowledgments

This work was primarily funded by the European Union (ERC-2021-STG, 4DPhytoHybrid, 101042148). Additional support was provided by the Knut and Alice Wallenberg Foundation via the KAW Project FATE, the Wallenberg Initiative Materials Science for Sustainability and the Wallenberg Wood Science Center (KAW 2018.0452.). Funding was also provided by the Swedish Government Strategic Research Area in Materials Science on Advanced Functional Materials at Linköping University (Faculty Grant SFO-Mat-LiU No. 2009-00971).

The authors acknowledge the facilities and technical assistance of the Umea Center for Electron Microscopy (UCEM) at the Chemical Biological Centre (KBC), Umeå University, a part of the National Microscopy Infrastructure NMI (VR-RFI 2019-00217). The authors wish to thank Agnieszka Ziolkowska (Umeå University) for the TEM imaging.

D.M. thanks Ayuda RYC2021-031668-I and PID2023-152287OA-100 financiada por MCIN/AEI/10.13039/501100011033 y por la Unión Europea NextGenerationEU/PRTR.

C.M. acknowledges support from the Swedish Research Council (2023-03651) and the Swedish Government Strategic Research Areas in Materials Science on Functional Materials at Linköping University (Faculty Grant SFOMatLiU no. 2009-00971). SV and LB acknowledge additional support from Åforsk Foundation (20-502).

## Author contributions

ES conceived the project. EC performed most experiments and data analysis. CM performed AFM experiments and data analysis. LB, XJ, AS and SV provided cells for starting cell cultures and shared expertise and guidance on cell culture. DP assisted with tensile test and mechanical characterization of auxetic and honeycomb ELMs and performed analysis of tensile data. AGF performed photosynthetic measurements and analysis. GD assisted with ETE-S polymerization and enzymatic assay. EM assisted with 3D bioprinting and maintenance of ELMs. DM synthesized ETE-S molecule. ES and EC designed the research. ES supervised the work. ES and EC wrote the manuscript with input from all authors.

